# Computational Model of Heterogeneity in Melanoma: Designing Therapies and Predicting Outcomes

**DOI:** 10.1101/2022.01.21.477232

**Authors:** Arran Hodgkinson, Dumitru Trucu, Matthieu Lacroix, Laurent Le Cam, Ovidiu Radulescu

## Abstract

Cutaneous melanoma is a highly invasive tumor and, despite the development of recent therapies, most patients with advanced metastatic melanoma have a poor clinical outcome. The most frequent mutations in melanoma affect the BRAF oncogene, a protein kinase of the MAPK signaling pathway. Therapies targeting both BRAF and MEK are effective for only 50% of patients and, almost systematically, generate drug resistance. Genetic and non-genetic mechanisms associated with the strong heterogeneity and plasticity of melanoma cells have been suggested to favor drug resistance but are still poorly understood. Recently, we have introduced a novel mathematical formalism allowing the representation of the relation between tumor heterogeneity and drug resistance and proposed several models for the development of resistance of melanoma treated with BRAF/MEK inhibitors. In this paper, we further investigate this relationship by using a new computational model that copes with multiple cell states identified by single cell mRNA sequencing data in melanoma treated with BRAF/MEK inhibitors. We use this model to predict the outcome of different therapeutic strategies. The reference therapy, referred to as “continuous” consists in applying one drug (or several drugs) without disruption. In “combination therapy”, several drugs are used sequentially. In “adaptive therapy” drug application is interrupted when the tumor size is below a lower threshold and resumed when the size goes over an upper threshold. We show that, counter-intuitively, the optimal protocol in combination therapy of BRAF/MEK inhibitors with a hypothetical drug targeting cell states that develop later during the tumor response to kinase inhibitors, is to treat first with this hypothetical drug. Also, even though there is little difference in the timing of emergence of the resistance between continuous and adaptive therapies, the spatial distribution of the different melanoma subpopulations is more zonated in the case of adaptive therapy.

## 1 Introduction

More than one half of melanomas carry mutations of the gene coding BRAF kinase, which is part of the MAPK signaling pathway, involved in cell growth and proliferation. In this pathway, BRAF phosphorylates and activates MEK that in turn phosphorylates and activates ERK, a potent effector that induces the transcription of many important genes that play dominant role in survival and development of tumour cells. In melanoma, targeted therapies based on BRAF inhibitors (vemurafenib, dabrafenib, encorafenib) and MEK inhibitors (trametinib, cobimetinib, binimetinib) aim at reducing the activity of this key signaling cascade [Gross et al.(2015), Zhang et al.(2015), McClure et al.(2021), Zhang and Bollag(2021)]. BRAF inhibitors act differentially on cancer and healthy cells. Indeed, elevated MEK and ERK activity is induced mainly by BRAF dimers, and less by monomers. In BRAF-mutated melanoma, RAS-GTP levels are insufficient to promote BRAF dimerization, therefore the inhibition of BRAF monomers is sufficient for ERK inactivation. This specificity reduces the toxicity of this type of treatment [Poulikakos and Rosen(2011)]. Although treatment based on these kinase inhibitors initially leads to efficient tumour regression, resistance appears almost systematically. Several mechanisms have been associated to acquired resistance, such as RAS mutation, receptor tyrosine kinase activation that either compromise ERK inactivation or induce other survival pathways such as PI3K/AKT. [Poulikakos and Rosen(2011)].

We focus here on a non-exclusive, but different cause of resistance, that involves the development of several drug tolerant cell states by non-genetic mechanisms. The non-genetic nature of adaptive resistance in melanoma is first suggested by the reversibility of this process: resistant tumors can re-sensitize upon a drug holiday [Das Thakur et al.(2013), Sun et al.(2014)]. Coexistence of sensitive and resistant cells with anti-correlated fitness in treated and untreated conditions can also explain apparent tumour re-sensitization in the absence of drug by positive selection of sensitive cells and negative selection of resistant cells, without the need for transitions between different cell states [Hodgkinson et al.(2019)]. Moreover, single cell RNA analysis has demonstrated plastic transitions between distinct cellular phenotypes in cell lines [Fallahi-Sichani et al.(2017), Smalley et al.(2019), Wouters et al.(2020)] and in primary derived xenograft (PDX) models [Rambow et al.(2018), Marine et al.(2020)] submitted to BRAF inhibitor therapy. The treatment induced transitions between states have robust features, common to many patient derived cultures and different cell lines [Wouters et al.(2020)]. Between the melanocytic and mensenchymal-like states which represent the sensitive and resistant extremes there are intermediate states resembling nutrient-starved cells and evolving via several trajectories towards mensenchymal-like states. The intermediate states and the trajectories originating therein show intrinsic variability of the gene expression, which suggests that the transitions between states are continuous rather than discrete [Fallahi-Sichani et al.(2017), Rambow et al.(2018), Wouters et al.(2020)].

These fundamental findings could be used to design new therapeutic strategies to avoid resistance. The re-sensitization, either real or apparent, arising when resistant cells are slowly growing in untreated conditions, suggest that a discontinuous adaptive treatment, alternating ‘on’ and ‘off’ drug periods, may be able to control tumor size, at least for some time. Combination therapy may also depend on one’s capacity to predict the changes induced by the primary tumour treatment, in space and time. For instance, drug tolerant neural crest stem cells, which are enriched upon treatment with BRAF/MEK inhibitors, display a RXR-driven signature, suggesting that these cells could eventually be targeted pharmacologically by using RXR-inhibitors [?]. Besides anti-BRAF targeted therapies, the recent discovery that immune checkpoint inhibitors, targeting regulatory molecules on T lymphocytes (anti-CTLA4, anti-PD-1, and anti-PD-L1), are highly efficient in melanoma patients has revolutionalized the treatment of metastatic melanoma. However, each treatment modality has limitations. While treatment with targeted therapies is associated with a strong beneficial short-term response but is followed by systematic resistance, treatment with immune checkpoint inhibitors has a lower response rate but associates with better long-term responses on a subset of melanoma patients. Thus, despite these considerable improvements in melanoma treatment, the development of new clinical strategies remains necessary and a better understanding of melanoma biology is likely to provide additional therapeutic options to patients with resistant cancers [Naderi-Azad and Sullivan(2020), Ma et al.(2021)].

In this paper we use a computational framework to study the heterogeneity of melanoma and develop a predictive model for various therapeutic outcomes.

## 2 Results

### 2.1 Multidimensional, data driven model of heterogeneity

Contrary to more traditional models of heterogeneity that consider a finite number of discrete cell states and transitions between these [Delitala and Lorenzi(2011)], our model can cope with a continuous spectrum of states. In this model, cell populations are represented as distributions (heatmaps) over many dimensions; spatial but also structural, representing internal cell-state variables such as gene expression, signaling, and metabolic activities (Figure 1a,b). An interesting possibility is to use single cell gene expression and feature extraction methods such as t-distributed stochastic neighbor embedding (t-SNE) in order to define reduced structural dimensions (Figure 1c,d). In this case, the distributions (heatmaps) predicted by the model (Figure 1b) can be directly compared to the empirical single cell distributions. We call this approach ‘mesoscopic’ as it is intermediate between a microscopic approach, which simulates each cell individually, and a macroscopic one, in which the cell-to-cell variability is averaged out.

**Figure 1:**
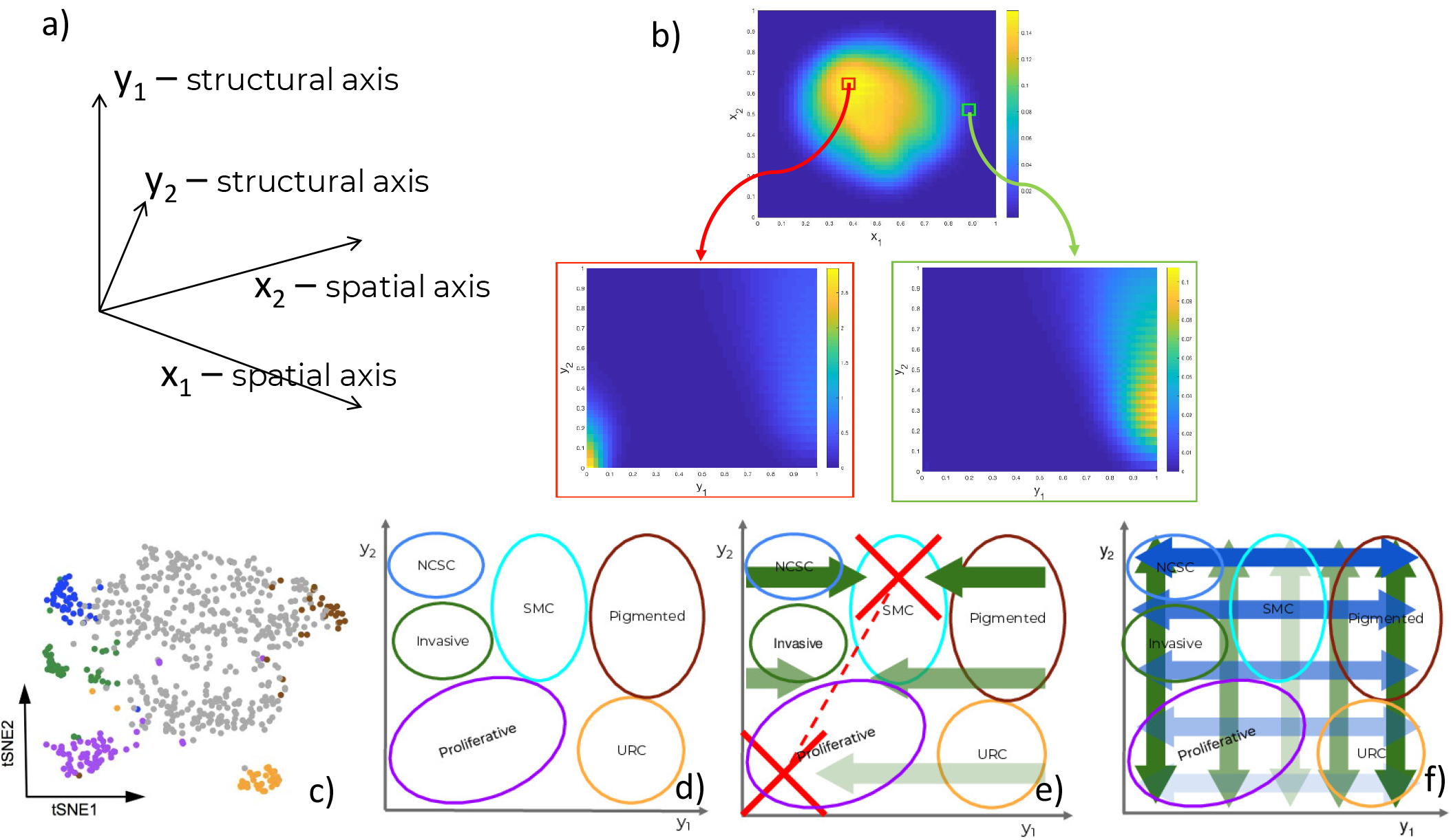
Components of the data driven heterogeneity model: a) Dimensions of the model. b) Multidimensional cell distributions predicted by the model emphasizing treatment induced zoning. c) Reduced representation of single cell expression data, from [?]. d) Cell states represented as domains in structural/gene expression space. e) Directed (advective) structural fluxes. f) Undirected (diffusive) structural fluxes.

In order to clarify the structural dimensions of our tumour, we have identified 6 different tumour subpopulations; namely proliferative, invasive, pigmented cells, neural-crest stem cells (NCSCs), starved-like melanoma cells (SMCs), and uncharacterised resistant cells (URCs). These are schematically represented in the structural plane in Figure 1d and are in line with previous experimental results [Rambow et al.(2018)]. It should be recalled that individual cells may concurrently occupy several states, existing in a continuum of gene expression across the structural domain.

The model predicts dynamic heterogeneity, meaning that the multidimensional distributions depend also on time. The evolution of these distributions are driven by spatial fluxes, involving undirected (diffusive) and directed (advective) spatial cell motility, and by structural fluxes, corresponding to changes of the cell state. The undirected structural fluxes (structural diffusion) correspond to random changes of the cell state leading to the spread of the cell distributions (increased heterogeneity) without changes of modal positions in the structural dimensions. The directed structural fluxes (structural advection) correspond to deterministic changes of the cell state, leading to shifts of the distribution modes. The cell distribution dynamics, represented as one 4D (2 spatial and 2 structural dimensions) partial differential equation (PDE), is coupled to five other 2D PDEs coping with the spatial distributions of other variables such as extracellular nutritional environment (ECNE), chemo-attractant (surrogate for mediated cell-cell communications), and drug concentrations. The effect of the drugs on the cells’ distribution is taken into account in the negative (degradative) source terms that depend on their position within the structural domain, i.e. on the cell state. For details, see Methods.

### 2.2 Targeted treatment exacerbates heterogeneity

The model recapitulates the dynamics of the cell heterogeneity observed in [Rambow et al.(2018)] (see Movies S1 & S2). Starting with a naïve tumor containing a population of sensitive melanocytes, several cell subpopulations are induced by the therapy. In our model, this is seen by the multimodality of the cell population’s structural distribution, with positions of the modes depending on time. As shown in Figure 1d, each sub-population is characterized by the position of the mode and by its spread in the structural domain.

The model predicts the typical three phase tumor growth curve under kinase inhibitors; a first phase wherein the tumour responds and shrinks, a second phase wherein the tumour is no longer visible corresponding to the minimal residual disease (MRD), and a third phase during which growth resumes corresponding to the development of resistance. During the MRD phase, heterogeneity strongly increases through continuous spreading of the cell distributions in the structural dimensions and, moreover, by development of co-existing, drug-tolerant, intermediate states between sensitivity and resistance (multi-modality, see Movie S2).

### 2.3 Order in combination therapy matters

We have tested, in our computational setting, combination therapies by successively applying two differing types of treatments: (1) using BRAF/MEK inhibitors (BRAF/MEKi) as in [Rambow et al.(2018)], and (2) a hypothetical cancer treatment (HCT). We have considered that the tumor has the same intrinsic dynamics, defined by the same diffusion and advection terms, for the two treatments. In particular, the cell states and their transitions will be the same for the two treatments. However, the two treatments eliminate cells differently, depending on their states. This difference between treatments was modeled by using a drug response function, defining how the drug dependent cell degradation changes with the cell state. This function peaks in the modal position of the primary tumour, in the case of BRAF/MEKi, or in the positions of the BRAF/MEKi resistant states, typically invasive and URC cell populations, in the case of HCT (see Methods and Supplementary Figure 1). Applied alone, the BRAF/MEKi treatment induces immediate and drastic tumour reduction, followed by MRD and development of resistance after approximately 4 months. The HCT treatment leads to a mild response initially, but like BRAF/MEKi treatment, induces tumor adaptation. However, the representation of invasive and URC cell states is only moderate because they are now more effectively eliminated.

Treatments using BRAF/MEKi (Figure 2a & Movie S3) or HCT (Figure 2b & Movie S4), alone, resulted in a re-establishment of initial tumour volume, prior to the end of the study period, with HCT inducing resistance far earlier than BRAF/MEKi. For the combination therapy, BRAF/MEKi then HCT, we observe a later time-point for the re-establishment of the initial tumour volume, in comparison to BRAF/MEKi only, but still resulted in a significant increased tumour growth rate (Figure 2c & Movie S5). Starting first with HCT and then using BRAF/MEKi, however, was a better strategy that significantly increases the delay to development of resistance and also reduces the tumor load by combining the advantages of the two treatments (Figure 2d & Movie S6).

**Figure 2:**
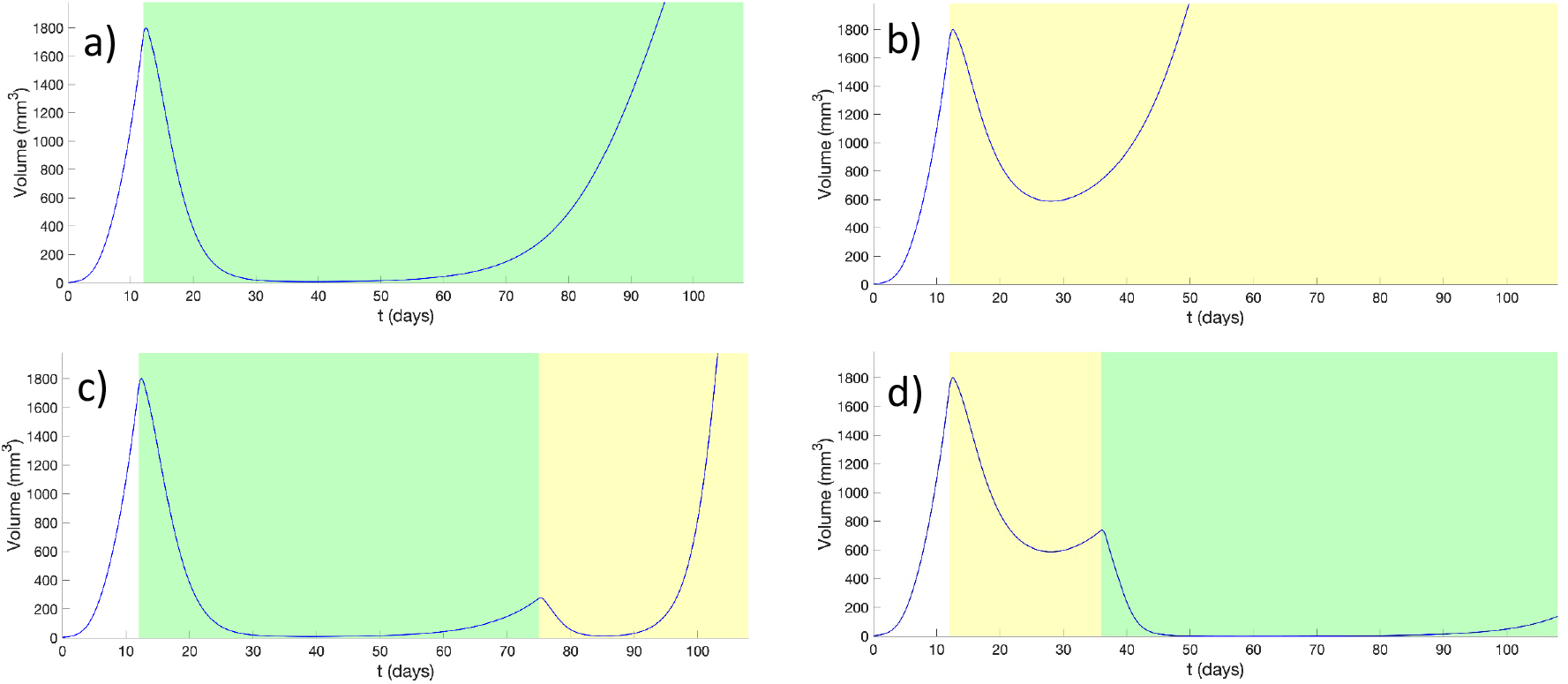
Outcomes from combination therapy, with treatment intervals indicated by graphical shading for BRAF/MEKi (*green*) and HCT (*yellow*). Panels show the outcomes from a) continuous BRAF/MEKi, b) continuous HCT, c) combination BRAF/MEKi then HCT, and d) combination HCT then BRAF/MEKi treatment regimes.

### 2.4 Output in terms of heterogeneity depends on the therapeutic strategy

The dynamics of melanoma cells submitted to kinase inhibitors is typically robust. In the case of adaptive therapy, although the intermediate dynamics is modified by allowing the tumour to grow before re-applying treatment, our model predicted that resistance development can not be avoided (Movies S7 & S8). However, in terms of spatial heterogeneity, the outcome is much more variable. In Figure 3 we have represented the spatial distribution of different cell states at the end of MRD and beginning of resistance, for various treatments. In all cases one can notice that cell states depend on position, a phenomenon called zoning. The details of this phenomena depend on the type of therapy. Adaptive therapy generates more pronounced zoning, with steeper and mutually exclusive patterns (Figure 3) than continuous therapies.

**Figure 3:**
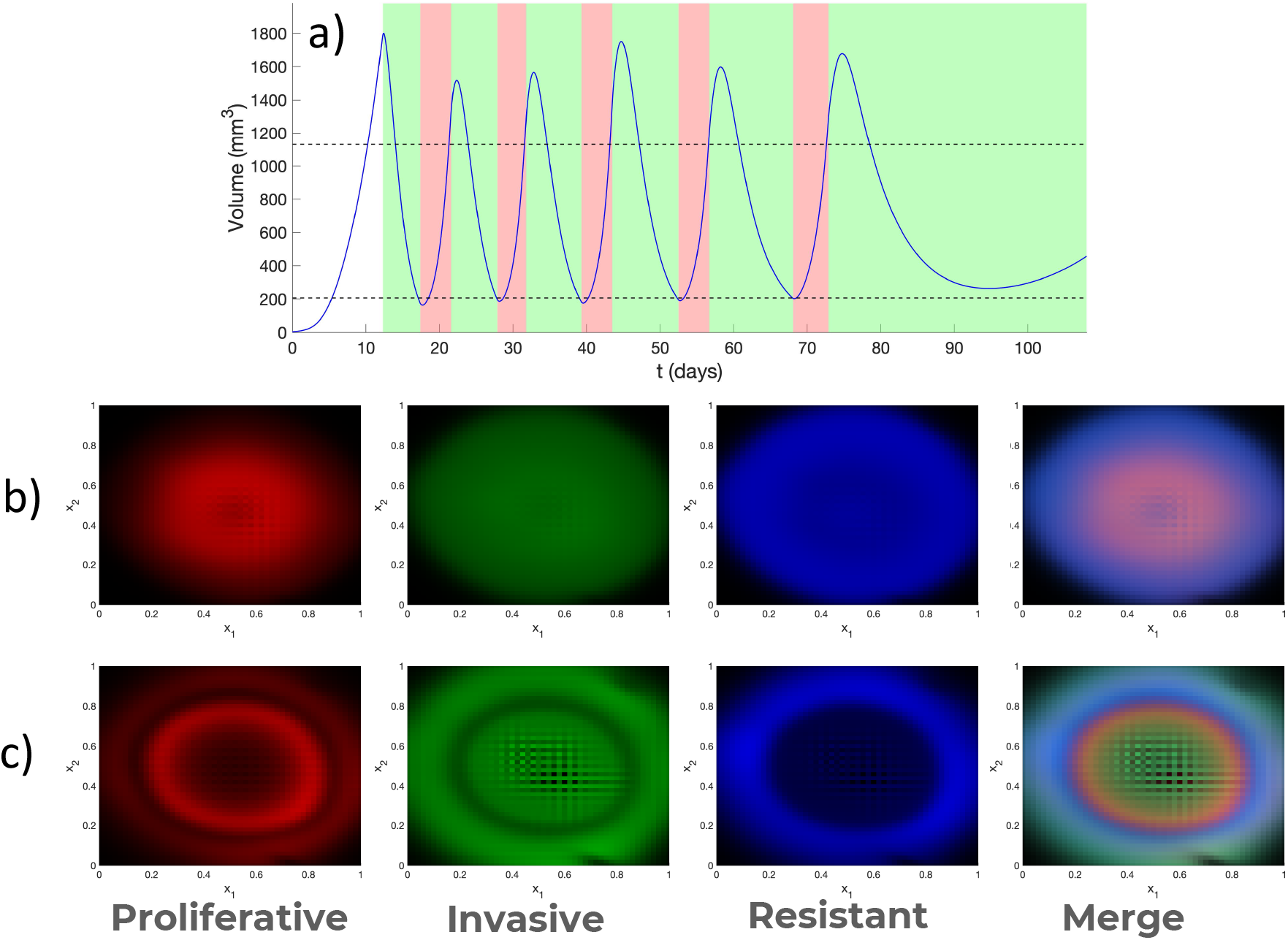
Adaptive therapy. a) Outcome of adaptive therapy, with BRAF/MEKi treatment intervals indicated in *green* and drug hollidays in *pink*. Decision about treatment is taken every day. Treatment is applied if the volume is higher than the upper threshold, stopped if the volume is lower than the lower threshold (thresholds are indicated as dotted lines). Spatial heterogeneity (zoning) generated by continuous b) and adaptive c) therapy.

## 3 Discussion and Conclusion

Treatment by kinase inhibitors leads to an heterogeneity upsurge in melanoma. At least part of this heterogeneity is generated by non-genetic mechanisms and involves continuous modification of the gene expression program leading to state transitions. Our mathematical model captures the essential features of these transitions and explains the heterogeneous dynamics by diffusive and advective spatial and structural fluxes. Moreover, our model predicts *in silico* the outcomes of various therapies.

We showed that, considering combination therapies, it is better to treat first with a less effective hypothetical drug, targeting sub-populations that develop during tumour resistance phases, before treating with BRAF/MEKi. The explanation of this rather counter-intuitive result can be found in the cell population dynamics. The intermediate response of the tumour to any of the treatments results in increased heterogeneity by diffusive processes. If the first applied treatment is BRAF/MEKi, this acts mainly on cells belonging to early modes and there is a non-negligible probability that some cells escape treatment and become resistant. The same probability is very small when the first applied treatment is HCT that acts preferentially on cells belonging to late modes; the role of HCT initial application is to avoid starting the BRAF/MEKi treatment with some cells that are not sensitive. Then, the application of BRAF/MEKi kills practically all the remaining cells and resistance takes much longer time to develop (see Movies S5 & S6). One should note that, due to structural diffusion, any cell state can, in theory, give rise to all other cell states. Therefore, in order to confine cells to BRAF/MEKi-sensitive modes, the drug has to act on a large domain of cell states, not only on a single intermediate drug tolerant sub-population. This is difficult to perform using targeted therapies.

A possible candidate for the hypothetical cancer treatment (HCT) is the drug family of immune checkpoint inhibitors (ICIs). Although this treatment does not act directly on melanoma cells, it can have a differential indirect effect on melanoma sub-populations, and acts more generally than targeted treatments. Very recent Phase III trials combining kinase inhibitors and ICIs show that starting with ICIs leads to better results in terms of survival time and duration of response than starting with kinase inhibitors [Atkins et al.(2021)]. This is explained by our theory if checkpoint inhibitors induce effective prior elimination of resistant stage sub-populations. There are, however, other interpretations of the interplay between kinase inhibitors and immunotherapy. Obenauf et al [Obenauf et al.(2015)] showed that kinase inhibitors induce changes in the stroma and cell secretome and hypothesize changes of immune cell composition. Other authors suggested that treatment with BRAFi leads to favorable changes in the tumour microenvironment in synergy with immune checkpoint inhibitors (see [Naderi-Azad and Sullivan(2020)] for a review). Little is yet known on the interactions between the immune system and melanoma cell sub-populations. We hope that future experimental and modelling work in the field, will elucidate the mechanistic aspects of these interactions.

Simple models of adaptive therapy were based on the idea that, in the absence of drug, resistant cells grow more slowly than sensitive cells [Smalley et al.(2019)]. It is believed that this fitness advantage allows sensitive cells to recover at least partially during a drug holiday. Although this effect is present in our model, it is compensated by structural and spatial diffusion that lead to increased heterogeneity and the delay increase of the time to resistance is only moderate. The resulting tumour, however, depends on the type, continuous or adaptive, of treatment. Our data suggest that a tumor resulting from adaptive treatment has more pronounced zoning.

From a theoretical perspective, our model shows the interplay between directed and undirected structural fluxes for the development of plasticity and heterogeneity. Undirected fluxes correspond to diffusion and random changes of cell states. As well known in physics, or in neutral theory of molecular evolution, free diffusion can reach any state from any other state if one waits a time proportional to the square of the state change. In the presence of treatment, diffusion is not free and has to cross barriers generated by the drugs action. In this case, the escape transition time is exponential. The escape transition time and the proportion of escaping cells depend on the position, height and width of the barriers, which are different for different treatments. This dependence further explains why order matters in combination therapy and why heterogeneity may differ when employing adaptive strategies, since the barrier is time-transient. Another important theoretical aspect is the symmetry breaking induced by the treatment. Although a barrier can be crossed in both directions, the transition probability is asymmetric if one of the states is more stable than the other. This leads to the notion of metastable states hierarchy, in which states are distinguished by the time that cells spend in each one of these; this time can be very very long for highly stable states. In adaptive therapy one tries to stabilize one metastable state by alternating treatment and holiday periods. The success of this strategy depends on conditions that may be difficult to guarantee, especially in a multidimensional context and for a spatially heterogeneous drug distributions.

We should nevertheless emphasize that our model is mostly phenomenological with structural dimensions representing nonlinear functions of the gene expression data. As several findings point towards the role of BRAFi in metabolic remodelling [Corazao-Rozas et al.(2013)], it would be very useful to interpret the structure variables in terms of metabolism. This is possible within our formalism as metabolic ODE models (see for instance [Jia et al.(2019)]) are transposable into structural advection fluxes, where metabolic stochasticity or uncertainty would translate to diffusive fluxes. This possibility will be investigated in future work. Furthermore, the distribution of blood vessels that are sources of nutrition and drug compounds is an important variable for understanding zoning aspects of cancer adaptation to treatment (see also [Kumar et al.(2019)]). Like in [Kumar et al.(2019)], we expect that sensitive and resistant cells’ spatial distributions depend on the distance to these biological sources. Blood vessel distributions can be reconstructed from *ex vivo* tumor sections [Kiemen et al.(2020)] and we will use these distributions to increase the realism of future models.

## 4 Methods

### 4.1 General formalism

Mesoscale models of cancer heterogeneity are based on partial differential equations and can be generically obtained from the Liouville continuity equation [Hodgkinson et al.(2018a), Hodgkinson et al.(2018b), Hodgkinson et al.(2019)]. Let us consider that there are *n* types of cells. In this model cells are distinguished by two types of variables, a discrete one representing the type *i* ∈ {1,…, *n*} and a continuous one *y* = (*y*_1_,…, *y_m_*) representing the internal state (vector of concentrations of biochemical species, for instance). Then *c* = (*c*_1_,…, *c_n_*) represents a vector of cell distributions satisfying the equation

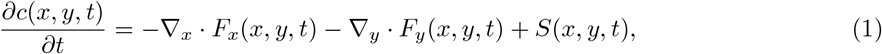

where *x* is the spatial position, *y* is the cell’s internal state (structure variable), *F_x_* is the spatial flux, *F_y_* is the structural flux and *S* is the source term. If the cell’s internal state follows ODEs 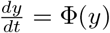, then the structural flux is advective *F_y_* = *c*Φ. If the cell’s state follows random Brownian motion in the structural space, then the structural flux is diffusive resulting from Fick’s law *F_y_* = –*D_y_*∇*_y_c*, where *D_y_* is the structural diffusion coefficient. The spatial flux contains terms related to cell motility: undirected (diffusion), or directed (chemotaxis, haptotaxis) [Chaplain and Lolas(2006)]. The source term integrates cell proliferation, death and transition from one cell type to another. It also copes with discrete stochastic changes (finite jumps) of the cell state *y*, other than those included in the continuous flux *F_y_*.

### 4.2 Model derived from single cell expression data

In this section we present only the broad lines of the model construction. The details can be found in the Supplementary Text.

#### Model components

Our melanoma progression model has two main components: the cancer cell population density *c*(*t, x, y*) and the the extra-cellular nutritional environment (ECNE) density *v*(*t, x*). In our minimal melanoma model space positions x and structural positions y are two dimensional (the latter correspond to the 2 axes of the t-SNE representation of the tumor transcriptome). We also consider spatial gradients of three types of diffusible molecules, namely 1) the nutritional molecular species, provided by the ECNE and consumed by cells, 2) the acidic molecular species, produced by cells and degrading the ECNE as in [Gatenby and Gawlinski(1996)], and 3) drugs.

The fluxes defining the model dynamics have been derived using the following assumptions:

#### Spatial variables and fluxes

We assume, consistently with previous mathematical studies of spatial cancer dynamics [Chaplain and Lolas(2006)], that the spatial dynamics of melanoma cells are governed by both random (diffusive) and deterministic (advective) components. The random (diffusive) component is assumed to occur as a result of tissue-scale reorientation and volume-filling processes. The deterministic (advective) component is assumed to result from directed cell motility and is driven by cell-environment interactions. In particular, we assume that cells exhibit controlled migration to sites of chemically elevated nutritional content (chemotaxis), as well as to sites of higher ECNE density (haptotaxis).

#### Structural variables and fluxes

The definition of the structural variables follows from the data analysis in [Rambow et al.(2018)]. Unsupervised clustering of single cell mRNA-seq data identified several types of cell sub-populations with distinct transcription states. The high-dimensional transcriptome was compressed to a 2D map using t-distributed stochastic neighbor embedding (t-SNE). The support of this 2D map is our structure space domain. The different transcription states represent sub-domains in this representation (see Figure1d). The cell-state transitions observed experimentally can be represented as diffusion and advective flow in this domain. The flow changes the positions of the cells in the 2D structure domain, moving them from one state to another. Thus, rather than considering a number of distinct cell types, we have build a model with only one cell type whose state can change continuously by the structural fluxes. A cell is added to a sub-population if its state enters the corresponding stuctural sub-domain and is subtracted if it dies or if it leaves the sub-domain. In order to define the structural fluxes, we start by identifying the sub-domains corresponding to different sub-populations inside the tumor at different times. Although seven transcriptional signatures were identified (Table S1 of [Rambow et al.(2018)]), we focus upon the description of 6 primary states important for resistance. For their localization in the structural domain we use cardinal points, as follows:

i. Melanoma cells with a “proliferative” signature are predominant in naive tumors, localized south-west (SW).
ii. Invasive cells are also present in naive tumors. They are localized east (E).
iii. Pigmented cells expressing markers of differentiation are induced by the treatment. They are localized north-west (NW).
iv. Neural crest stem cells (NCSC) are enriched by the treatment, have a maximum during the minimal residual disease and are diluted out during the development of resistance. They are localized north-east (NE).
v. Starved-like melanoma cells (SMC) are rapidly induced by the treatment, and become predominant during MRD. They are localized north (N).
vi. Uncharacterised resistant cells (URCs) were not thoroughly biologically investigated, though the model predicts they may have a biological interest. They are localized south-west (SW).

The structural fluxes describe the metabolic adaptation within the structural domain and diffusion-like exchanges between cell populations (Figure1e & f). In order to define these fluxes we use the following dynamical assumptions:

- Horizontal advection is assumed to stabilise the proliferative (SW) state, since there is no known emergence of URCs prior to resistance;
- SMC states (N) are also stabilized by horizontal advection fluxes that converge towards this state, allowing cells to populate this minimally mitotic state;
- advection is assumed to interpolate linearly at intermediate phenotypes between proliferative cells and SMCs;
- horizontal diffusion is assumed to be maximal in the northern regions of the structural plane and decrease towards southern regions, characterising a low exchange rate between proliferative and URC states;
- vertical diffusion is maximal towards the western and eastern regions – allowing exchange between proliferative, invasive, and NCSC or pigmented and URC populations – but lower exchange rates between SMC and southern states.

In principle, by diffusion any cell state can give rise to all cell states. However, advection maintain a certain degree of cellular hierarchy. These assumptions have been made upon a reasoned analysis of the figures presented in [Rambow et al.(2018)], as a minimal set of functional assumptions to reproduce observed patterns, but do not necessarily represent an optimal or biologically motivated set of assumptions.

#### Source terms and degradation

The source and degradation terms describe cell proliferation and death, respectively. We use the (known) metabolic activity [Rambow et al.(2018)] as a proxy for proliferation; thus, mitosis is significantly reduced among SMC cells and increased among proliferative cells. We consider that treatment is the only cause of active cell death. Due to the nature of our modelling framework, drugs may target cells with a spectrum of specific expression markers as would be the case in the clinical scenario. In this case, we assume the existence of two particular treatments. Firstly, BRAF and MEK inhibitors were employed within the study conducted by Rambow *et al*. [Rambow et al.(2018)] and, as such, are assumed to primarily target a distribution centred around the proliferative population, stretching into the invasive population but with diminished success among cells in the NW of the structural domain (Supplementary Figure 1). Secondly, a hypothetical cancer therapeutic (HCT) has been used for the sake of illustration and targets primarily the invasive and URC cell populations, with an expansive effectiveness span E and SW (Supplementary Figure 1).

#### Spatial dynamics of other components

Given the complexity of the dynamics in the primary cancer cell population, the dynamics of other components have been kept as simple as possible. It is assumed that the ECNE exhibits only a natural restorative growth process, as well as acidic species-induced and natural degradation kinetics, with respective rate constants. Nutritional and acidic species exhibit diffusion, as well as controlled production, and natural degradation. Finally, the drug species also exhibit diffusion, time dependent administration, as well as natural and cell-based degradation.

## Author Contributions

AH and OR conceived the project. OR wrote the paper with help from AH, ML and LLC. AH designed the model, numerical scheme, and performed the simulations. DT contributed to the general formalism and numerical scheme.

## Acknowledgments

The authors would like to thank Jean-Christophe Marine for very helpful discussion.

## Supplemental Data

Supplementary Text: Detailed model description (includes Supplemental Table 1)

Supplementary Figure 1: Structural drug effectiveness function for BRAF/MEK inhibitor a) and for HCT drug b). The drug acts mainly on cells whose states are located at maxima of the effectiveness function.

Supplementary Movie 1: Spatial (x-) distribution under continuous BRAF/MEK inhibitor treatment.

Supplementary Movie 2: Structural (y-) distribution under continuous BRAF/MEK inhibitor treatment.

Supplementary Movie 3: Drug induced heterogeneity under continuous BRAF/MEK inhibitor treatment. The false colours of the tumour identify proliferative states (*red*); NCSCs, SMCs, invasive, and pigmented cells (*green*); and URCs (*blue*).

Supplementary Movie 4: Drug induced heterogeneity under continuous HCT treatment. The false colours of the tumour identify proliferative states (red); NCSCs, SMCs, invasive, and pigmented cells (*green*); and URCs (*blue*).

Supplementary Movie 5: Drug induced heterogeneity under combination therapy; first BRAF/MEK inhibitors then HCT. The false colours of the tumour identify proliferative states (red); NCSCs, SMCs, invasive, and pigmented cells (*green*); and URCs (*blue*).

Supplementary Movie 6: Drug induced heterogeneity under combination therapy; first HCT then BRAF/MEK inhibitors. The false colours of the tumour identify proliferative states (*red*); NCSCs, SMCs, invasive, and pigmented cells (*green*); and URCs (*blue*).

Supplementary Movie 7: Drug induced heterogeneity under adaptive therapy. The false colours of the tumour identify proliferative states (*red*); NCSCs, SMCs, invasive, and pigmented cells (*green*); and URCs (*blue*).

Supplementary Movie 8: Structural (y-) distribution of cancer cells under continuous BRAF/MEK inhibitor treatment.

The code is accessible at https://github.com/oradules/Melanoma2D_2021/.

## Supplementary Methods

### Description of the Mathematical Model

#### Cancer cell population dynamics

Using a novel framework (given in eq. (1) of the main text) for higher-dimensional mathematical modelling of population dynamics in cancer [1], we are here able to represent the dynamics of the population in time, space, and approximated gene expression levels. The approach results in partial differential equations in a number of dimensions equal to the spatial dimension plus the gene expression dimensions. The latter are reduced to 2 through the use of t-distributed stochastic neighbourhood embedding (t-SNE) method on single cell transcriptomics data [2].

The cell population is represented by a density function *c*(*x, y*) defined on spatial and structural variables (*x, y*), where *x, y* ∈ [0, 1]^2^. The spatial density is the marginal density *c_x_*(*x*) = *∫ c*(*x, y*)*d_y_*.

Cell population evolves as a result of spatial fluxes, structural fluxes, and cell sources (proliferation and degradation).

Spatial fluxes occur when cells move in space. As in previous models of the spatial evolution of cancer [3, 1], our model considers that cells may move either in a disorganised, or in an directed manner. The disorganised movement corresponds to the space diffusion flux that, according to the Fick law, is proportional to the gradient of cell density and is directed opposite to this gradient. Directed movement corresponds to chemotactic, or haptotactic fluxes that are proportional to chemical gradients, or to the gradient of ECNE, respectively. In both cases cells move uphill gradients, towards higher densities of cells in chemotaxis, or towards higher density of ECNE in haptotaxis. We consider two types of chemotactic gradients: *m*_1_ (*x*) is a nutrient produced by ECNE and *m*_2_(*x*) a chemo-attractant produced by the cancer cells. The spatial flux term is, therefore, given by:

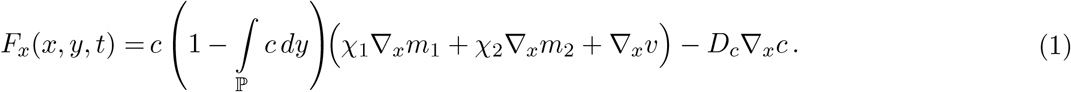

The factor 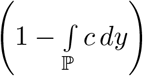 has a stabilizing role and was introduced to avoid the Keller-Siegel instability [4].

Structural fluxes occur when cells change their states. In contrast to spatial fluxes that result from thermodynamics, spatial fluxes are derived from observations on the single cell data. Structural fluxes are also of two types, diffusive and advective, corresponding to random, zero mean, changes of the cell state and deterministic changes of the cell state, respectively. For the diffusive terms, we assume that these vary with the two structural variables *y*_1_ and *y*_2_ as described in the Methods section and Figure 1f in the main text. The advective terms are similarly described in Methods and Figure 1e in the main text and are assumed to have the proliferative and SMC subpopulations, as well as all the points linearly interpolated between these states as stable steady states. The stability of these states was considered higher for larger *y*_2_, which was modelled as a linear factor (1 – *r_min_*)*y*_2_ + *r_min_*, 0 < *r_min_* < 1, in the advection flux. Thus, we have used the following model for structural fluxes:

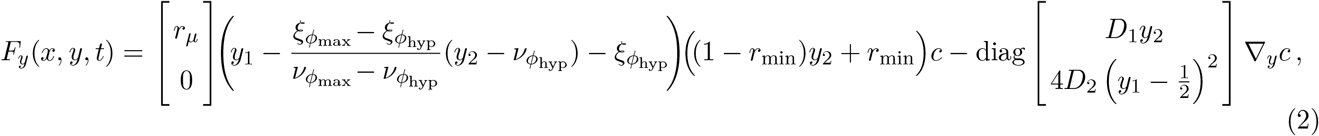

where (*ξ*_*ϕ*hyp_, *ν*_*ϕ*hyp_), (*ξ*_*ϕ*max_, *ν*_*ϕ*max_) are the structural space centroids of the SMC and the proliferative cells subpopulations, respectively.

Cell sources terms are of two types: positive proliferation terms, and negative degradation terms.

Degradation terms are proportional to a combination of the drug concentrations *p*_1_*f*_1_(*y*) + *p*_2_*f*_2_(*y*), where *p*_1_, *p*_2_ are the drug concentrations and *f*_1_(*y*), *f*_2_(*y*) are effectiveness functions of the two drugs (see Figure S1). The drug effectiveness functions cope with the fact that cells with different states *y* are eliminated differently by the two drugs.

The proliferation terms take into account competition on resources (logistic growth) and are proportional to the concentration of nutrient *m_1_* produced by ECNE. The proliferation term also contain a factor depending on the cell state *y*, that mimics the cell metabolic activity derived in [2]. For modelling this factor, it is assumed that all cells have a basic proliferation rate, *ϕ_c_ϕ*_0_ that there is a region (centered in (*ξ*_*ϕ*up_, *ν*_*ϕ*up_), in between the states defined in the main text, and close to the west part of the structural domain) exhibiting elevated proliferative activities and an elevated rate, *ϕ_c_ϕ*_up_; and that the cells in the proliferative state exhibit the greatest proliferation rate, *ϕ_c_ϕ*_max_. Meanwhile, the SMC state is assumed to exhibit a significantly lower rate than the remainder of the domain and this is achieved by dividing the entire term by 1, plus a Gaussian function centered in (*ξ*_*ϕ*hyp_, *ν*_*ϕ*hyp_). We use the following expression for the source terms:

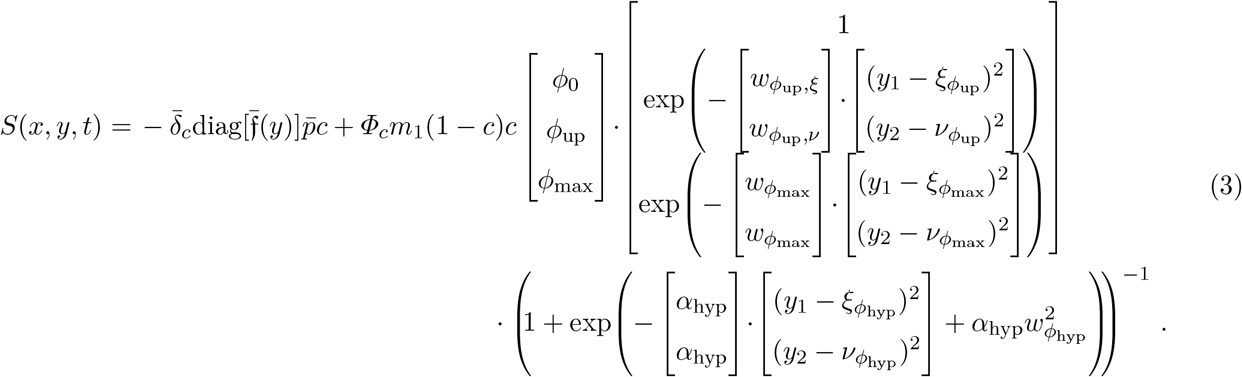

#### Spatial dynamics of other components

As well as the cancer cell population, three types of other species are considered in our model; namely the extra-cellular nutritional environment (ECNE), the chemical species, and the drug species.

As in several previous models of cancer invasion [5, 3, 6, 1], we assume that the cancer cell population invades the stroma by degrading the ECNE to make space for invading cells. Hence, there is a chemically-mediated degradation, with the rate vector 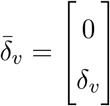 (only the acidic species *m*_2_ degrade ECNE). The degradation of the *δ*_*v*,0_ ECNE is also achieved through a natural decay term, with rate *δ*_*v*,0_. Furthermore, the ECNE grows logistically with a rate *ϕ_v_*.

The chemical species, themselves, diffuse with a rate vector 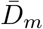. In particular, the current model considers only two chemical species; the nutritional species *m*_1_, produced by the ECNE, and the acidic or degradative species *m*_2_, produced by the cancer cells. The production of these species is given by the rate vector 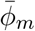. Logistic factors *m_i_*(1 – *m_i_*), *i* ∈ {1, 2} multiply the production terms, in order to keep the chemical species concentrations bounded 0 ≤ *m_i_* ≤ 1 for *i* ∈ {1, 2}. Chemical species are assumed to undergo degradation with rates given by the vector 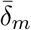.

We consider two drug species, representing BRAF/MEKi (for simplicity we consider only one species even if two kinase inhibitors are administered) and the hypothetical cancer treatment (HCT), respectively. The drug species exhibit diffusive spatial dynamics, with a rate vector 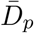. The two drug species, are differentially administered to the patient, on the basis of the required treatment. This differential treatment is mathematically represented by the time-dependent, user-defined, function, 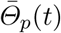. Drug species are assumed to be degraded by natural processes, with rates 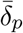, and by cellular uptake and degradation, with rates 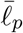.

#### Full System of Equations

Combining the above equations with the assumptions for the ECNE, chemical, and drug populations, we may then write the following complete set of PDEs as:

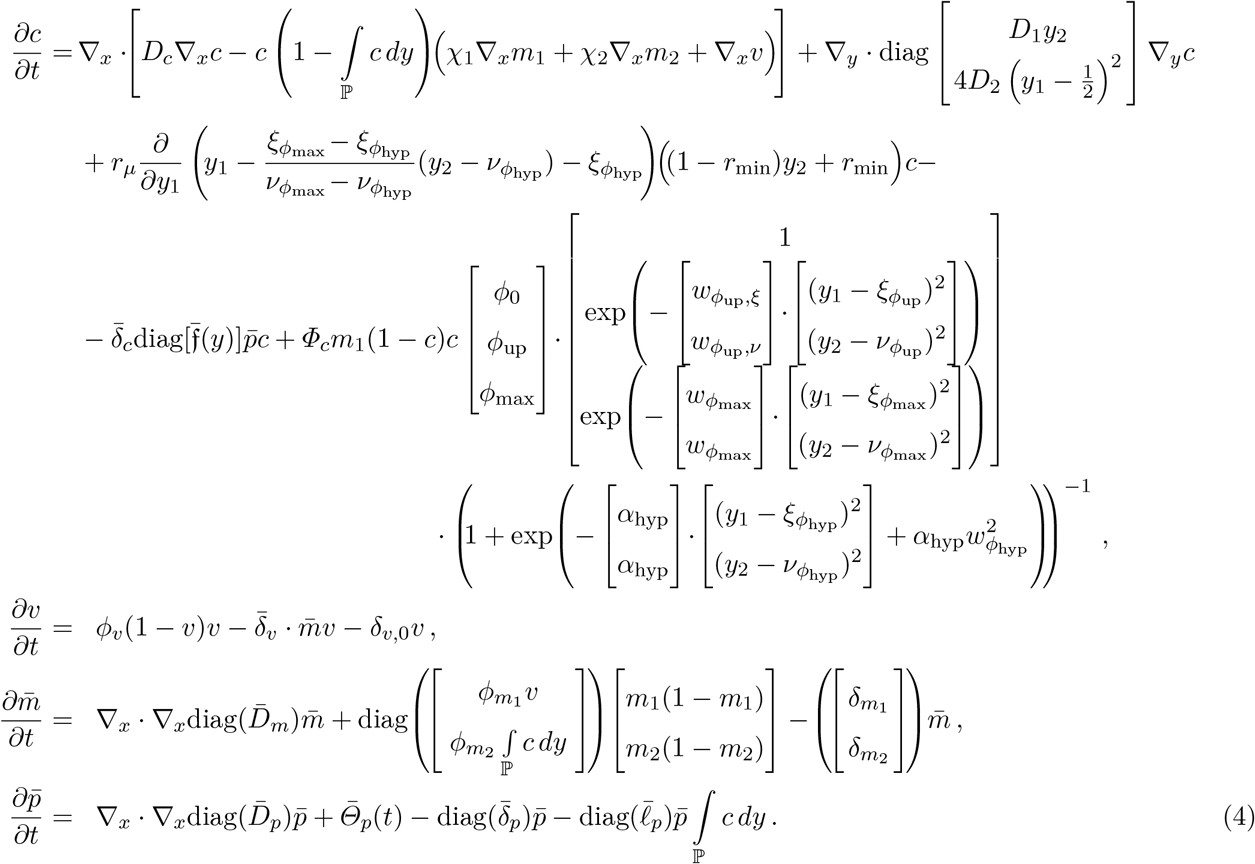

The appropriate biological interpretation and value of the model parameters are provided in the Table 1.

**Table 1:** Model variables and parameters. The units are arbitrary.

| Parameter | Interpretation | Value | Variable | Interpretation |
| --- | --- | --- | --- | --- |
| $(D_1, D_2)$ | Structural diffusion coefficients | $(10^{-3}, 2.10^{-3})$ | $c$ | cancer cell density |
| $\bar{D}_c$ | Spatial diffusion coefficient | $[10^{-5}, 10^{-5}]^T$ | | |
| $(\chi_m, \chi_v)$ | Chemo- and Hapto-taxis coeff. | $(2.10^{-4}, 2.10^{-4})$ | | |
| $r_\mu$ | Structural advection coefficient | $12.10^{-3}$ | | |
| $r_{min}$ | Vertical structural position | 0.17 | | |
| $\phi_c$ | Basic proliferation rate | 0.5 | | |
| $\phi_0$ | Relative proliferation rate | 0.08 | | |
| $\phi_{up}$ | Relative proliferation rate | 0.45 | | |
| $\phi_{max}$ | Relative proliferation rate | 0.47 | | |
| $(\xi_{\phi_{up}}, \nu_{\phi_{up}})$ | Centre of elevated proliferation | (0.5, 0.2) | | |
| $(\xi_{\phi_{max}}, \nu_{\phi_{max}})$ | Centre of maximum proliferation | (0.15, 0.15) | | |
| $(\xi_{\phi_{hyp}}, \nu_{\phi_{hyp}})$ | Centre of hypometabolism | (0.5, 0.8) | | |
| $(w_{\phi_{up}, \xi}, w_{\phi_{up}, \nu})$ | Width of elevated proliferation | (2, 35) | | |
| $w_{max}$ | Width of maximum proliferation | 20 | | |
| $(\alpha_{hyp}, w_{hyp})$ | Width of hypometabolic region | (100, 0.25) | | |
| $\bar{\delta}_c$ | Drug induced cell death rate | $[0.5, 0.5]^T$ | | |
| $\phi_v$ | ECM regeneration | 0.02 | $v$ | ECNE: extra-cellular<br>nutritional environment |
| $\delta_v$ | $\bar{m}$ dependent ECNE degradation | 0.05 | | |
| $\delta_{v0}$ | Basal ECNE degradation | 0.01 | | |
| $(D_{m_1}, D_{m_2})$ | Chemical diffusion coefficients | $(2.10^{-4}, 2.10^{-4})$ | $m_1$ | Nutritional molecules |
| $(\delta_{m_1}, \delta_{m_2})$ | Degradation rates $m_1, m_2$ | (0.2, 0.02) | $m_2$ | Degrading molecules |
| $(\phi_{m_1}, \phi_{m_2})$ | Production rates $m_1, m_2$ | (0.1, 0.1) | | |
| $(D_{p_1}, D_{p_2})$ | Drug diffusion coefficients | $(1.2.10^{-3}, 1.2.10^{-3})$ | $p_1$ | BRAF/MEKi |
| $(\delta_{p_1}, \delta_{p_2})$ | $c$ dependent degradation | (0.65, 0.65) | $p_2$ | Hypothetical cancer |
| $(\delta_{p_1,0}, \delta_{p_2,0})$ | Basal drug degradation | $(5.10^{-3}, 5.10^{-3})$ | | treatment (HCT) |

The code is accessible at https://github.com/oradules/Melanoma2D_2021/.

**Figure S1:**
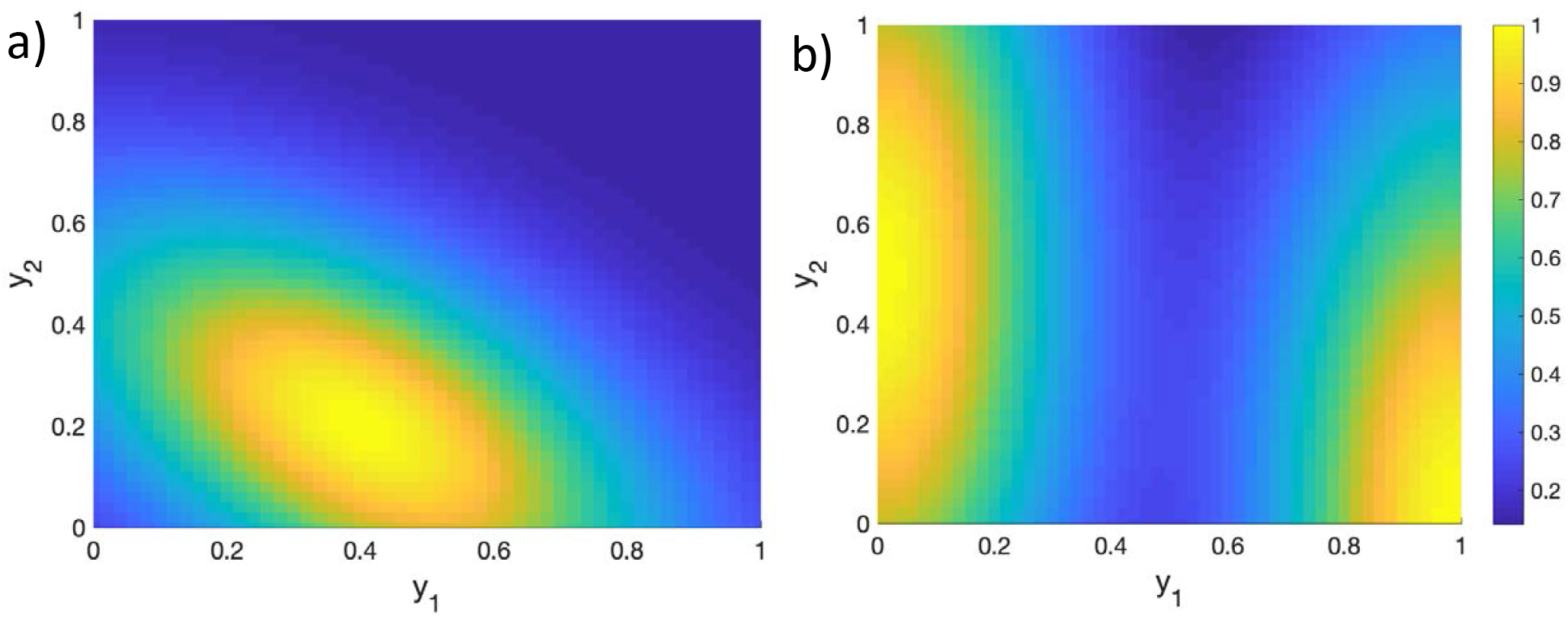
Structural drug effectiveness function for BRAF/MEK inhibitor a) and for HCT drug b). The drug acts mainly on cells whose states are located at maxima of the effectiveness function.

