## Supplementary figures and images for "Computational Model of Heterogeneity in Melanoma: Designing Therapies and Predicting Outcomes"

### Supp Figure 1

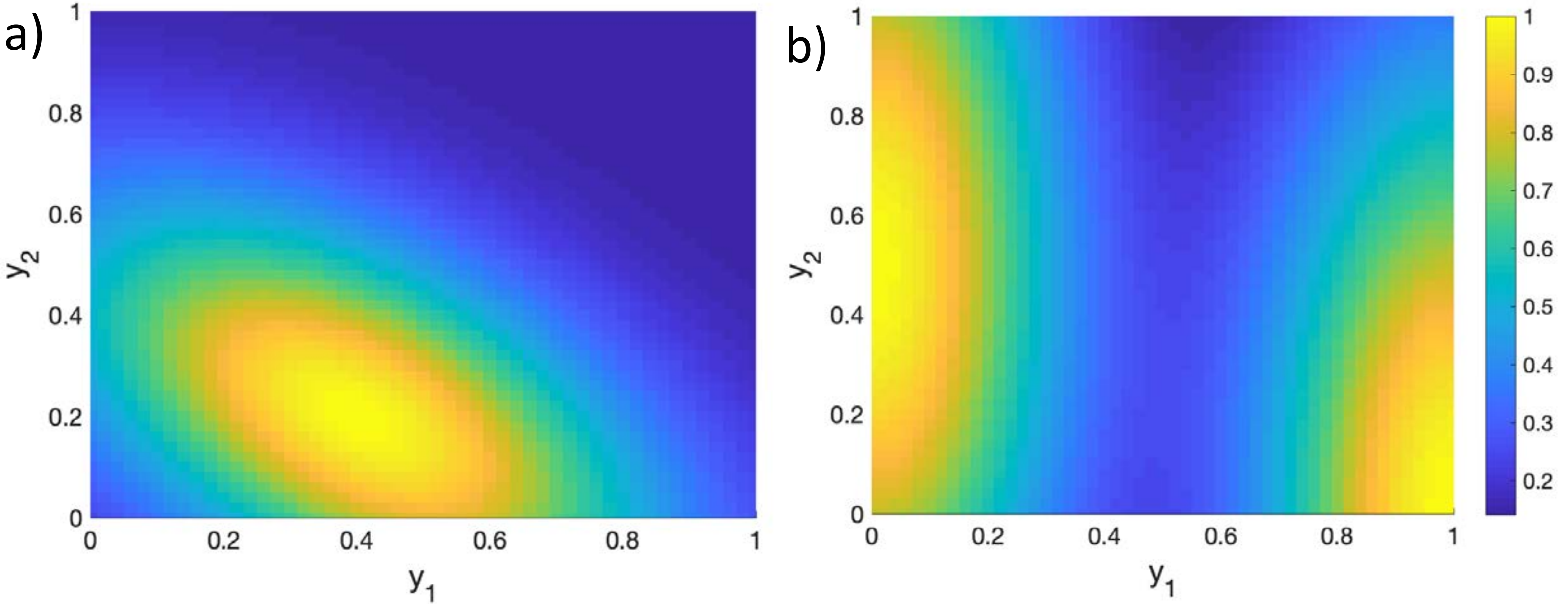
